# ARA: accurate, reliable and active histopathological image classification framework with Bayesian deep learning

**DOI:** 10.1101/658138

**Authors:** Łukasz Rączkowski, Marcin Możejko, Joanna Zambonelli, Ewa Szczurek

## Abstract

Machine learning algorithms hold the promise to effectively automate the analysis of histopathological images that are routinely generated in clinical practice. Any machine learning method used in the clinical diagnostic process has to be extremely accurate and, ideally, provide a measure of uncertainty for its predictions. Such accurate and reliable classifiers need enough labelled data for training, which requires time-consuming and costly manual annotation by pathologists. Thus, it is critical to minimise the amount of data needed to reach the desired accuracy by maximising the efficiency of training. We propose an accurate, reliable and active (ARA) image classification framework and introduce a new Bayesian Convolutional Neural Network (ARA-CNN) for classifying histopathological images of colorectal cancer. The model achieves exceptional classification accuracy, outperforming other models trained on the same dataset. The network outputs an uncertainty measurement for each tested image. We show that uncertainty measures can be used to detect mislabelled training samples and can be employed in an efficient active learning workflow. Using a variational dropout-based entropy measure of uncertainty in the workflow speeds up the learning process by roughly 45%. Finally, we utilise our model to segment whole-slide images of colorectal tissue and compute segmentation-based spatial statistics.

## Introduction

Histopathological images of cancer tissue samples are routinely inspected by pathologists for cancer type identification and prognosis. Hematoxylin-Eosin (H&E) stained slides have been used by pathologists for over a hundred years. With such long history and proven applicability, histopathological imaging is expected to stay in common clinical practice in the coming years^1^. With the advent of digital pathology, histopathological images became available for automated analysis at scale^2^. To this end, a rich catalogue of machine learning approaches to image classification and whole-slide segmentation has been developed^3, 4^, promising to aid the effort of pathologists in interpreting the images^5^. Such machine learning models need to be perfectly *accurate*, as classification errors may result in faulty disease diagnosis and patient treatment. On top of that, we stipulate that in application to digital pathology, the models should also be *reliable* in their predictions. When performing the difficult task of automated classification or diagnosis based on histopathological images, they should state uncertainty in their predictions, indicating difficult cases for which human expert inspection is necessary. While accuracy is optimised by every machine learning method, reliability is another desired feature that is not delivered by many state of the art solutions.

Recent years brought particularly intensive development of deep-learning based approaches to image classification. In particular, Convolutional Neural Networks (CNNs) have served as a backbone for numerous breakthroughs in computer vision as a whole, specifically in image classification. Since 2012, when the groundbreaking AlexNet was created by Alex Krizevsky^6^, the state of the art has rapidly shifted from machine learning algorithms using manual feature engineering (henceforth referred to as ‘traditional’ machine learning approaches) to new deep learning ones^7^. Medical imaging in general^8–12^, and histopathological image classification in particular^13–20^, became important applications of these methods. Multiple machine learning methods go beyond the tasks of tissue type classification and whole-slide segmentation, confirming there is more information about the patients encrypted in histopathological images than immediately visible by eye^5^. For example, *Wang et al.*^21^ trained a CNN to predict survival of patients from their pathological images. Another deep learning model^22^ allowed prediction of mutations in several genes from non–small cell lung cancer histopathology. Finally, Bayesian measures of measuring uncertainty for deep-learning methods have been proposed^23, 24^ and successfully applied to medical image analysis^25^. Those developments open the avenue to reliable image classification, where prediction uncertainty can be reported together with the predicted class.

All of these exciting methodological inventions would not be possible without training data that is numerous enough to train accurate models. Publicly available training data with annotated images such as the Breast Cancer Histopathological Database^26^ allow algorithm benchmarking and evaluation, sparkling new method developments^27, 28^. Similarly, *Kather et al.*^29^ released a colorectal cancer dataset with H&E tissue slides, which were cut into 5000 small tiles (or patches), each of them annotated with one of eight tissue classes. They also devised an efficient classification method, with image-derived features serving as basis for a support vector machine model. Since its publication, this dataset was utilised several times to verify the performance of an array of methods. *Ribeiro et al.*^30^ developed a traditional method that uses multidimensional fractal techniques, curvelet transforms and Haralick descriptors. They tested its accuracy using the *Kather et al.* dataset in a binary classification scenario. *Wang et al.*^31^ developed a Bilinear CNN architecture that takes as input H&E stained images decomposed into H and E channels and used all eight classes from the *Kather et al.* dataset to verify its performance. *Pham*^32^ utilised this dataset to assess their autoencoder architecture. *Sarkar et al.*^33^ created a new saliency-based dictionary learning method and used the *Kather et al.* dataset for both training and testing. Finally, *Ciompi et al.*^34^ used it as an independent test set for an evaluation of two stain normalisation strategies. All traditional methods reported accuracy lower than the original classification method by *Kather et al.*. The AUC value in the eight-class classification task obtained by *Wang et al.*^31^ was higher than the one achieved by *Kather et al.*, confirming that a CNN is the method of choice for this dataset as well. Notably, none of these methods aimed for reliability, as they did not assess the uncertainty of their predictions.

Generation of datasets like the ones described above requires laborious workload of pathologists who process whole-slide images and assign labels to selected image regions. The requirement of meticulous pathological annotation limits the amount of data available for model training. Formally, this relates to the bias-variance trade-off in machine learning^35^. The effort to minimise bias on a small training set may result in high variance and low accuracy on unseen data, an effect known as overfitting. In order to minimise the expected test error, model regularisation techniques penalising model complexity can be applied. For CNNs, a technique called dropout has been proposed as a means of regularisation^36^. Another technique, called active learning, can be used to deal with the difficulty of laborious data annotation. Active learning is an iterative procedure, where in each step the model is re-trained on data expanded with new samples, which are added based on results from the previous steps in order to maximise the learning rate. Variants of active learning depend on the way the new samples are selected. One method of choice is selection which maximises the diversity of the training set^37^. *Gal et al.*^38^ proposed an active learning procedure for Bayesian deep learning models, where new samples are added in each iteration based on their uncertainty estimated using variational dropout. This technique can be conceptualised by an analogy to a diligent student, who while taking a course actively asks the teacher for more examples on topics which are hard for them to understand. There are several attempts at active learning for histopathological image classification in the literature, using both traditional machine learning^37, 39–42^ and deep learning^43–46^. However, none of these approaches utilised uncertainty for selection of new samples in active training.

In this work, we introduce an accurate, reliable and active (shortly, ARA) learning framework for classification of histopathological images of colorectal cancer. To this end, we develop a new CNN model (called ARA-CNN) for classification of colorectal cancer tissues, trained on the *Kather et al.* dataset (Fig. 1). The model achieves stellar accuracy, higher than reported in the original publication of *Kather et al.*^29^ and in later studies. The key contribution of this work is an extensive analysis of the utility of two variational-dropout based uncertainty measures, Entropy *H* and BALD, in their application to histopathological image classification. We demonstrate that the distribution of uncertainty is increased for tissue classes that are the most difficult to learn for the model. Moreover, images that are mis-classified tend to have the highest uncertainties. We propose an active learning framework, where the model suggests the most uncertain classes for annotation by a pathologist (Fig. 1A). We show that *H* outperforms random selection of images and BALD when applied to select samples in an active learning procedure, speeding up the learning process by roughly 45%. In-depth inspection indicates that images with very low Entropy *H* are highly characteristic of each tissue class. On the contrary, images with very high *H* are atypical or show pathological features that could be shared by other classes, which makes them pathologically difficult to categorise. In addition to image classification and uncertainty estimation, the framework is successfully applied to image segmentation and provides segmentation-based statistics of tissue class abundance in whole tissue slides (Fig. 1B).

**Figure 1.**
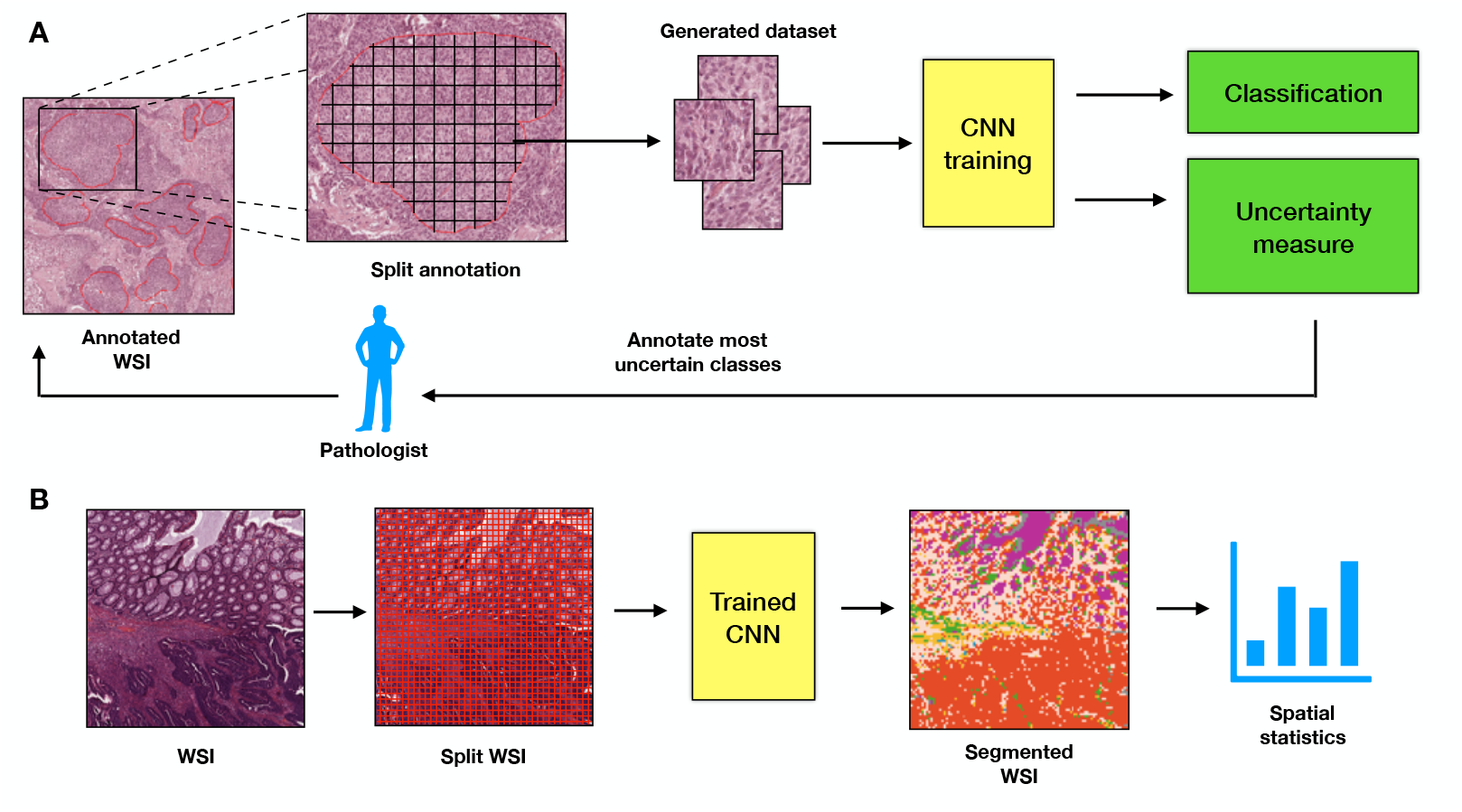
Overview of the proposed ARA framework. **A** Active histopathology workflow. Annotated whole-slide images (WSIs) are split into small image patches, which constitute a dataset. ARA-CNN is trained on that dataset. After the first round of training, the pathologist should be informed which classes are the most uncertain and prioritise them in annotating whole-slide images for the next round. We then take new annotated whole-slide images and continue the workflow until we reach a satisfying level of classification accuracy. **B** Segmentation workflow. Whole slide images are split into small image patches. Each of these is classified by trained ARA-CNN and is assigned a colour based on its classification result. These coloured tiles are merged together to form a segmented whole slide image and can be analysed in terms of their spatial relationships. Each resulting tile has a measured uncertainty value as well, so pathologists can make an informed decision whether to take the automated segmentation as-is or to inspect it manually.

## Methods

### Analysed data

The analysed dataset holds 5000 image patches belonging to eight balanced classes of histopathologically recognisable tissues^29^. The patches were pulled from ten anonymised and digitised tissue slides, stained with the H&E technique. After initial coarse-grained annotation, 625 non-overlapping tiles were extracted from contiguous tissue areas for each class. Each tile has the same size of 150 × 150 pixels (equivalent to 74*µm* × 74*µm*). The eight tissue classes are: tumour epithelium, simple stroma (homogeneous composition, includes tumour stroma, extra-tumoural stroma and smooth muscle), complex stroma (containing single tumour cells and/or few immune cells), immune cells (including immune-cell conglomerates and sub-mucosal lymphoid follicles), debris (including necrosis, haemorrhage and mucus), normal mucosal glands, adipose tissue, background (no tissue). Here, for the sake of brevity, these classes are labelled as: Tumour, Stroma, Complex, Lympho, Debris, Mucosa, Adipose and Empty. In addition, one tissue slide denoted by *Kather et al.*^29^ as a test image was used for the purpose of segmentation (see below).

### ARA-CNN model

To automatically classify the images from the analysed dataset into their corresponding classes, we developed and trained a Convolutional Neural Network (CNN) model. The architecture of the model, called ARA-CNN, was inspired by many state of the art solutions, including Microsoft ResNet^47^ and DarkNet 19^48^. For normalisation and to reduce overfitting, we used a popular technique called Batch Normalisation^49^. In ARA-CNN, overfitting is also reduced by using dropout^36^. This in turn allowed us to apply variational dropout^50^ during testing. For every tested image, the model provides not only its predicted class, but also a measure of uncertainty estimated using variational dropout.

### Network architecture

The ARA-CNN network accepts RGB images of size (128, 128, 3) as its input (Fig. 2A), where the values represent respectively: vertical resolution, horizontal resolution and the number of colour channels. The images from the training dataset were downsized to these dimensions. Input values are propagated to the first part of the network called stem (Fig. 2B). The stem contains a convolutional layer consisting of 64 filters, with filter size of (7,7) and stride size of (4,4). This is directly followed by max pooling with window size of (2,2), and identically sized strides. The output from this part is of size (16,16,64), where the values are: reduced width, reduced height and the number of filters. These operations decrease the spatial dimensions by a factor of 8, which in turn significantly reduces memory usage and can be considered an adaptation of network topology to a relatively simple texture structure of the input^51^.

**Figure 2.**
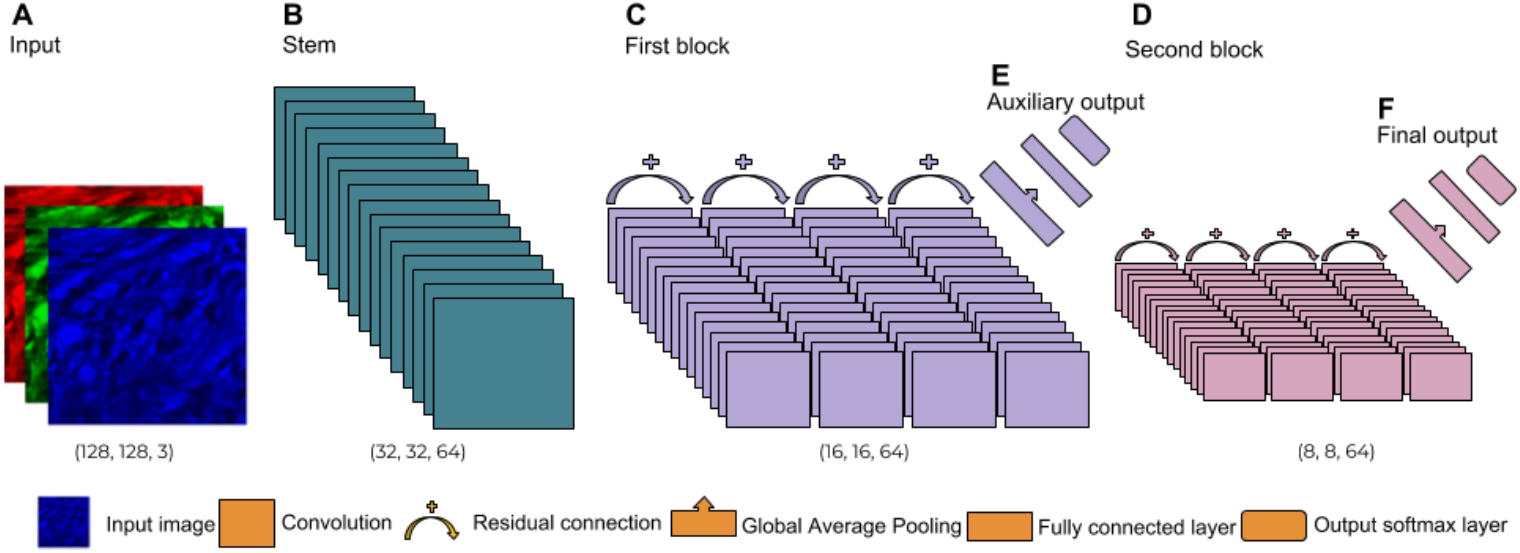
Structure of the ARA-CNN model. The network takes as input RGB images with dimensions of 128×128 pixels. They are passed to the stem, which contains a convolutional layer responsible for reducing the spatial dimensions of the input. This is followed by the first block and the second block, responsible for learning low-level and high-level image features respectively. Both of these blocks consist of four residual sections, with each of these sections containing a convolutional layer and a residual connection. The model has two outputs overall - an auxiliary output from the first block and a final output from the second block. Both of them use the softmax activation function.

The stem is followed by the first block (Fig. 2C). The main aim of this part is to learn and extract initial discriminative low-level image features. It consists of 4 residual sections, where the input to each block is transformed by a convolutional layer with 64 filters - each sized (3,3) and with stride of size (1,1). The result of this convolution is added back to the input, which creates a residual connection. The final section of this block is followed by an average pooling with window size of (2,2). This makes the output of this part of the network to be shaped (8,8,64). The next part of the model is the second block (Fig. 2D), which learns and extracts the final discriminative features - this time they are more high-level and abstract. Its structure is the same as that of the first block. After the final average pooling with window size (2,2), the output from this part is of size (4,4, 64).

The model has two outputs in total: the auxiliary output (Fig. 2E) and the main output (Fig. 2F). The main purposes of the former are to provide a better training signal to the stem and the first block (by making the features more discriminative) and to deal with the vanishing gradient problem^52^ during training. The output from the first block is transformed by global average pooling. Next, it is transformed by a fully-connected layer with 32 filters and dropout with rate of 0.5 (explained below in *Dropout*). Finally, it is fed to the fully-connected output layer with a softmax activation function. The final output is used for making the actual predictions. After the second block, the data is transformed by exactly the same set of transformations as in the auxiliary output - global average pooling, then a fully-connected layer with dropout, followed by a final output layer.

If not stated otherwise, each convolutional filter has dilation and stride set to 1. Additionally, each layer in the network (except the outputs) connects to a Batch Normalization layer. When deep learning models are trained, the distribution of inputs for each layer changes as a result of modified parameters in preceding layers, which slows down the whole process. Batch Normalisation combats this by normalising layer inputs for each training mini-batch. This enables the use of higher learning rates and significantly speeds up the training. Batch Normalisation also acts as a regulariser and reduces overfitting. The activation function used throughout the model is Leaky ReLU^53^:

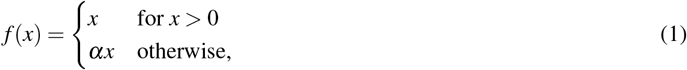

where the parameter *a* is set to 0.1 and *x* is a weighted sum of inputs to a network unit.

### Dropout

Due to their size, deep learning models are especially prone to overfitting - they can inadvertently learn from sampling noise instead of actual non-linearities in the training data. One of the more popular and successful methods of combating this problem is dropout^36^. It works on the basis of randomly removing units in a neural network during training in order to simulate a committee of multiple different architectures. In our model, dropout is applied to two fully connected layers with 32 units preceding auxiliary and final output. Its rate is equal to 0.5, which means that during both inference and training approximately half of all units are turned off and set to 0.

### Model training

The whole dataset of 5000 images was split into a training dataset and a test dataset used for evaluation. Their sizes varied depending on the experiment. In the 8-class case, we randomly divided the dataset into the training set with 4496 images (562 images per class) and the test set with 504 images (63 images per class). In the case of binary classification, the training set contained 1124 images, while the test set was comprised of 126 images (divided in half between Tumour and Stroma). These divisions were repeated ten times in the process of 10-fold cross-validation. Moreover, we also performed 5-fold cross-validation and 2-fold cross-validation, with the dataset split according to the number of folds.

Additionally, in each training epoch the training data was split into two datasets: the actual training data and a validation dataset. The latter was used for informing the learning rate reducer - we monitored the accuracy on the validation set and if it stopped improving, the learning rate was reduced by a factor of 0.1. In the task of evaluating model performance, this split was in proportion 90% to 10% between actual training data and the validation set, respectively, while in active learning the split was in proportion 70% to 30%. This is due to the fact that in active learning we start from a very small dataset and 10% was too small of a proportion to provide enough validation samples.

For parameter optimisation, we used the Adam^54^ optimiser. The training time differed depending on the experiment. In the cross-validation experiments we used 200 epochs, but in the active learning experiments it was 100 epochs instead (due to limited computational resources). In all cases, the training data was passed to the network in batches of 32, while the validation and test data was split into batches of 128 images.

### Loss function

During training, the categorical cross-entropy loss function was applied to both (auxiliary and final) outputs. The final loss is a weighted sum of these two losses with weight 0.9 for the final output and 0.1 for the auxiliary output. For observation *o*, a set of *M* classes and class *y*\* ∈ {1…*M*}, we denote the probability of assigning the observation to that class as 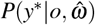, where 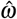 represents the estimated parameters of the model. Categorical cross-entropy can then be defined as:

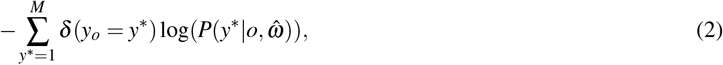

where *δ* is the Dirac function and *y*_*o*_ is the correct class for observation *o*.

### Variational dropout for inference and uncertainty estimation

In order to provide more accurate classification as well as uncertainty prediction, we adopted a popular method called variational dropout^50^. The central idea of this technique is to keep dropout enabled by performing multiple model calls during prediction. Thanks to the fact that different units are dropped across different model calls, it might be considered as Bayesian sampling from a variational distribution of models^24^. In a Bayesian setting, the parameters (i.e. weights) *ω* of a CNN model are treated as random variables. In variational inference, these random variables are posited to come from an approximating (variational) distribution *q*(*ω*). Thus, we assume that 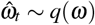, where 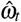 is an estimation of *ω* resulting from a variatonal dropout call *t*. Denote 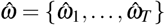. With these assumptions, the following approximations can be derived^24^:

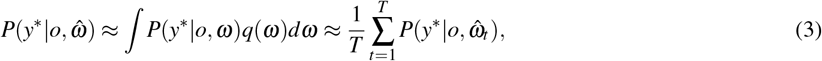

where *T* is the number of variational samples. In our model we used *T* = 50.

Variational dropout has enabled us to measure the uncertainty of predictions. We implemented two uncertainty measures: Entropy *H* and BALD^23^. If the output of the model is a conditional probability distribution 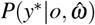, then the measure *H* can be defined as entropy of the predictive distribution:

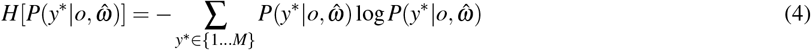

The second uncertainty measure, BALD, is based on mutual information and measures the information gain about the model parameters *ω* obtained from classifying observation *o* with label *y*\*. In the case of variational dropout, this can be expressed as the difference between entropy of the predictive distribution and the mean entropy of predictions across multiple model calls:

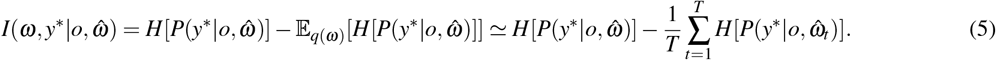

The difference between these two measures pertains to how they react to two different types of uncertainty in the data: epistemic and aleatoric^55^. The former type is caused by a lack of knowledge - in terms of machine learning, this is analogous to a lack of data, so the posterior probability over model parameters is broad. The latter uncertainty is a result of noise in the data - no matter how much data the model has seen, if there is inherent noise then the best possible prediction may be highly uncertain^23^. In general, the Entropy *H* measure cannot distinguish these two types of uncertainty. If uncertainty of a new observation is measured by *H*, then the value would not depend on the underlying uncertainty type. On the other hand, it is believed that BALD measures epistemic uncertainty of the model^23^, so it would not return a high value if there is only aleoratic uncertainty present. Depending on the dataset, one of these measures might work better than the other at catching and describing the uncertainty.

### Image segmentation

To perform segmentation of test tissue slides from the *Kather et al.*^29^ dataset, each of these 5000×5000px images was split into 10000 non-overlapping test samples with resolution of 50×50 pixels. These test images were then supplied as input to our model (by being upscaled to 128×128 pixels), which returned a classification into one of eight classes of colorectal tissue. Since the output of the model is a probability distribution, we selected the class with the highest value as the prediction for a given test image patch. We did not consider the measured uncertainty in this process. To get the final segmentation, we assigned a colour to each predicted class and generated a 50×50 pixels single-coloured patch for each test image. These patches were then stitched together to form the final images. Lastly, we applied a blurring Gaussian filter to smooth out the edges of tissue regions.

Finally, we performed a simple spatial analysis for each slide by counting the percentage of surface area taken by each class.

### Active learning

Active learning is an iterative procedure, where the initial model is trained on a small dataset and in consecutive iterations it is re-trained on a dataset extended by new samples. At each step, the new samples are added according to some acquisition function evaluated using the current model. Intuitively, the uncertainty measures described above are a good basis for an acquisition function in deep learning. In a given iteration, the model should first choose the samples it is most uncertain of^38^.

In this work we implemented and compared effectively three different acquisition functions. Two were based on uncertainty measures H and BALD, whereas the third was a random selection and served as a baseline. We performed a series of experiments in order to determine if uncertainty-based active learning can speed-up training with the colorectal cancer dataset. To this end, we emulated the proposed active learning workflow (Fig. 1A) utilising the available data. We started from generating three random splits of the full dataset - this gave us three test sets of 504 images and three training sets of 4496 images. Then for each of these test-train pairs, we performed the active learning procedure for both uncertainty measures plus a baseline training process based on random selection of images. In each case, we started from selecting 40 images per class (so 320 in total) from the training dataset. We trained the model on that small dataset and then, based on a given acquisition function, we chose 160 images to add to the previous 320. This slightly larger set became a new training dataset. We repeated this process, adding 160 images in each step, until there were no more images to draw from the initial full training dataset, giving us 28 training steps in total. Additionally, in order to eliminate the effects of random weight initialisation, we pre-initialised the model 8 times for each step and used these initialisations for each of the 3 dataset splits. Thus, for each of the 28 steps we had to train the model 24 times.

For the random selection, the 160 new images in each step were sampled uniformly at random from the full training dataset. For the uncertainty-based functions, we performed inference on remaining images from the full training set in each step. We evaluated the uncertainty for each image using the H and BALD measures, according to Eq. (4) and Eq. (5). We sorted the results by uncertainty in descending order and selected the top 160. The results for each active learning step were averaged between initialisations and dataset splits.

## Results

### Model performance

To evaluate the performance of ARA-CNN, similarly to previous models trained on the same dataset, we measured its receiver operating characteristic (ROC) curves, area under the ROC curves (AUC) and error rates in 10-fold cross-validation for both 8-class and 2-class (Tumour vs Stroma) classification tasks. In addition, we also evaluated precision-recall curves. We used images with all colour information preserved. The results were compared to those of the original model by *Kather et al.* (Fig. 3), as well as to other methods that used the same dataset. Where necessary, we performed 5-fold or 2-fold cross-validation and used the results as a comparison point. In their work, *Kather et al.*^29^ tested the performance of several low-level image features in combination with four classification algorithms, applied to grayscale images from their dataset. Their approach is an example of a ‘traditional’ procedure, where image features have to be hand-crafted and chosen appropriately depending on the dataset. The best results were reported for a combination of features containing: pixel value histograms, local binary patterns, gray-level co-occurence matrix and perception-like features. The best performing classifier was a support vector machine (SVM) algorithm with the radial basis function (RBF) kernel.

**Figure 3.**
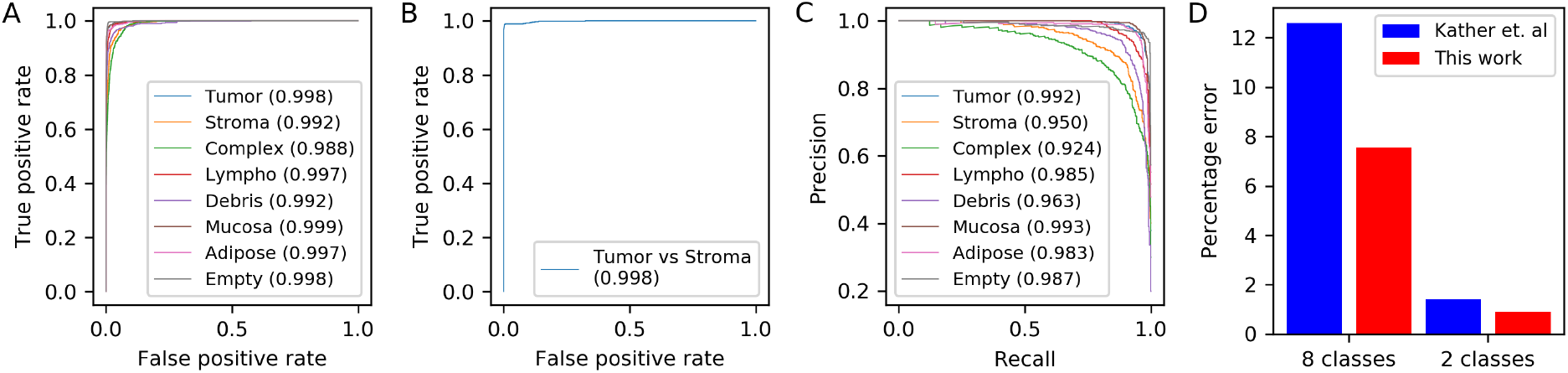
Model performance in 10-fold cross-validation. **A** ROC and area under the ROC curve (AUC) for classification into eight tissue types. The model presented in this work achieved an average AUC of 0.995 (a mean was taken across all eight classes), **B** ROC and AUC for binary classification between Tumour and Stroma. ARA-CNN achieves AUC of 0.998. **C** Precision-recall curves for ARA-CNN in a multiclass classification setting. The mean AUC for these curves is 0.972. **D** Error comparison to previous work. With error rate 7.56% for eight class classification, our model substantially reduces the error (by 5.04%) compared to error rate 12.6% of the best model assessed by *Kather et al*^29^. For binary (Tumour versus Stroma) classification, our model has error rate 0.89%, which is also lower than the 1.4% error rate of the Kather *et al.* model.

The ROC curves (generated with a one-vs-all method) for the 8-class experiment show excellent performance of ARA-CNN (Fig. 3A). The AUC values for the Tumour, Mucosa, Lympho, Adipose and Empty classes range from 0.997 to 0.999. Values for the Stroma, Complex and Debris classes are a little lower (from 0.988 to 0.992), which indicates that the model cannot always distinguish them from other classes. Still, the mean AUC value is 0.995, which is higher than the value of 0.976 obtained by *Kather et al*^29^. The ROC curve for the 2-class problem (Fig. 3B) and its corresponding AUC value of 0.998 also illustrate near-perfect performance of ARA-CNN. It is important to note that performance evaluation using ROC curves for the multiclass classification task in a one-vs-all setting may be biased due to the fact that the classes are unbalanced. In such a setting, it is better to use precision-recall curves (Figure 3C). The AUC values for these curves, as obtained by ARA-CNN, are a bit lower than for the ROC curves, but with the mean AUC of 0.972 are still indicative of excellent performance. The lowest AUC value (0.924) is obtained for the Complex versus all classification task. This indicates that the Complex class is the most difficult one to classify correctly for the model. We do not compare these results to other methods, as we are not aware of any other approaches that used precision-recall curves for performance evaluation on this dataset.

In terms of error rates, for the 8-class problem the ARA-CNN model reached an average rate of 7.56%, which is substantially lower, by 5.04%, than the best result reported by *Kather et al.*^29^ (Fig. 3D). Similarly, in the binary classification task, we obtained an error rate of 0.89%, lower than 1.4% obtained by *Kather et al*. Thus, our model is better than the best of standard approaches presented by *Kather et al.*^29^, especially in the multiclass classification scenario. One of the differences between deep learning and the standard approaches is that the former construct the features on the fly based on the data itself. Here, the features identified by ARA-CNN as part of the learning process outperform the set of features that were engineered by *Kather et al.*^29^ in the difficult task of decisively describing all classes in a multiclass image classification problem.

The classification performance of ARA-CNN is also superior or comparable to other published models that used the *Kather et al.* dataset, including both traditional and deep learning approaches that utilise CNNs (Table 1). ARA-CNN outperforms the traditional methods by a significant margin both in terms of AUC and accuracy. When it comes to CNN methods, *Wang et al.*^31^ performed 5-fold cross-validation and reported a mean AUC value of 0.985 (lower by 0.01 than ARA-CNN) and 92.6% accuracy (higher by 0.36% than ARA-CNN) for their BCNN in the multiclass task. Although BCNN and ARA-CNN achieve similarly high performance results, their architectures are very different. BCNN depends on an external method to perform stain decomposition of H&E images and is composed of two simple feed-forward CNNs, which take as input separate signals from the Eosin and Hematoxylin components and whose outputs are combined by bilinear pooling. We took a more typical deep-learning approach, with a deeper network with residual connections, where no independent feature extraction nor decomposition is needed, and the network itself is responsible for extracting important signals from raw image data. *Pham*^32^ used an autoencoder architecture to re-sample the images from the *Kather et al.* dataset and trained a small supervised network for different re-sampling factors. They reported at best an accuracy of 84.00% for binary classification, which is lower by 14.88% in comparison to our result. *Ciompi et al.*^34^ used the *Kather et al.* dataset for testing their model trained on an independent colorectal cancer dataset and reported relatively small accuracies of 50.96% and 75.55%, where the former was achieved without stain normalisation and the latter was an improvement resulting from having stain normalisation applied. However, since this model was trained on a different dataset, we do not directly compare our result to theirs. Overall, ARA-CNN’s achieves excellent performance on the *Kather et al.* dataset, and scores better than most other published methods that utilised the same data for training. Exceptional performance of our approach indicates that it successfully combines the flexibility typical for deep neural networks with strong regularisation resulting from dropout and Batch Normalisation.

**Table 1.** Comparison of different methods that used the *Kather et al.* dataset for training. ACC– accuracy. We summarise performance measures of compared methods as reported by the authors. Results in bold are the best in their category. * The authors do not explicitly state the number of folds. Since in other reported results the number of folds they used is 10, we assume 10-fold cross-validation here as well.

| Method | Method type | Problem type | Max. reported 10-fold ACC | Max. reported 5-fold ACC | Max. reported 2-fold ACC | 10-fold AUC | 5-fold AUC |
| --- | --- | --- | --- | --- | --- | --- | --- |
| *Kather et al.* | Traditional | Binary | 98.6% | - | - | - | - |
|  |  | Multiclass | 87.4% | - | - | 0.976 | - |
| *Ribeiro et al.\** | Traditional | Binary | 97.68% | - | - | - | - |
| *Sarkar et al.* | Traditional | Multiclass | 73.66% | - | - | - | - |
| *Wang et al.* | CNN | Multiclass | - | **92.6 ± 1.2%** | - | - | 0.985 |
| *Pham* | CNN | Binary | - | - | 84.00% | - | - |
| ARA-CNN | CNN | Binary | **99.11 ± 0.97%** | **98.88 ± 0.52%** | **98.88%** | **0.998** | **0.999** |
|  |  | Multiclass | **92.44 ± 0.81%** | 92.24 ± 0.82% | **88.92 ± 1.95%** | **0.995** | **0.995** |

### Tissue slide segmentation

In histological image analysis, the labelling of image patches is only the first step in the process of segmentation. To get a full overview of a tissue slide, it is necessary to see how image patches of different classes are placed in relation to each other and to measure their relative abundance. In particular, it is interesting to determine the neighbourhood of tumour cells. For example, the tumour being infiltrated by immune cells may be a marker of good prognosis. *Kather et al.*^29^ showed a simple segmentation approach using standard classification methods. We present a recreation of their procedure using the ARA-CNN model (see *Image segmentation*).

An example segmentation of a full tissue slide from the *Kather et al.*^29^ dataset is presented in Fig. 4. The segmentation can obviously be improved - the approach with stitching image patches is after all quite rudimentary. However, it can be good enough to see the aforementioned spatial relationships. As a basic spatial statistic, for each slide we generated a summary of tissue class distribution (Fig. 4). Histograms such as these can be used as a filter to find images for further consideration (e.g. those with high tumour concentration) in an automated diagnosis system.

**Figure 4.**
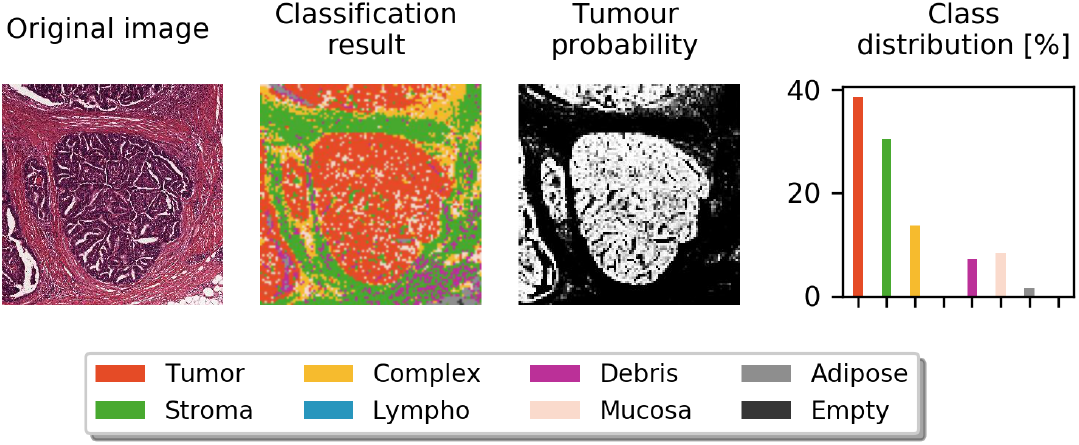
Segmentation. Segmentation of a single large tissue slide from the colorectal cancer dataset. The leftmost column presents the original WSI, the second one shows the segmentation done with our classification algorithm, while the third one is a visualisation of the Tumour class probability (the lighter the segment, the more probable that there is a tumour there). The last column contains a class distribution histogram - each bar represents the percentage of a given class in the segmented image.

### Uncertainty and active learning

Deep learning models are often criticised for being so-called black-boxes. Due to their complexity, it can be very hard to tell why a given test sample is classified to a certain class. The model presented in this work, thanks to its implementation of dropout and variational inference, has a few ways to measure the uncertainty of each prediction. These uncertainty measures allow the model predictions to be reliable. Consider an example image, which is classified by the model as Tumour with high probability 0.95, but the measured uncertainty is also high. This can mean that the prediction cannot be taken for granted and needs to be double-checked by a human. This additional indication of prediction uncertainty brings us one step closer to alleviating the problem of the black-box nature of deep learning and increases model-based understanding of the data. Here, we evaluated two uncertainty measures, Entropy *H* and BALD (see *Uncertainty estimation*), checking their distribution in each class and their performance as acquisition functions in active learning on the *Kather et al.*^29^ dataset.

First, we applied the trained model to 504 test images. For each image, we recorded the classification and the measured uncertainty. The results for Entropy *H* are presented in Fig. 5A. On average, the highest uncertainty values were reported for images from the Stroma and Complex classes. The biggest variance in uncertainty was measured for the Debris class. These three classes were also misclassified as each other, which indicates that they are similar in appearance and the model has a hard time differentiating them. This is in agreement with the precision-recall curves in Fig. 3D and with the analysis described below in *Understanding uncertainty*. In addition, it can be observed that misclassification occurred almost exclusively when the uncertainty was high. Thus, a high uncertainty is indeed a good indicator that the prediction may be faulty. The results for BALD are shown in Fig. 5B. On average, the most uncertain classes according to that measure are Stroma and Complex, in agreement with Entropy *H*. Interestingly, BALD measured much less variance in the Debris class, which makes Lympho the most variable class in this case. Moreover, the Empty class is relatively more certain according to BALD than in the Entropy *H* experiment. These differences may be a result of epistemic and aleatoric uncertainties present in the data, which are measured differently by BALD and Entropy *H* (see *Active learning*). Nevertheless, the BALD measure still captures the fact that misclassifications take place mainly for highly uncertain predictions.

**Figure 5.**
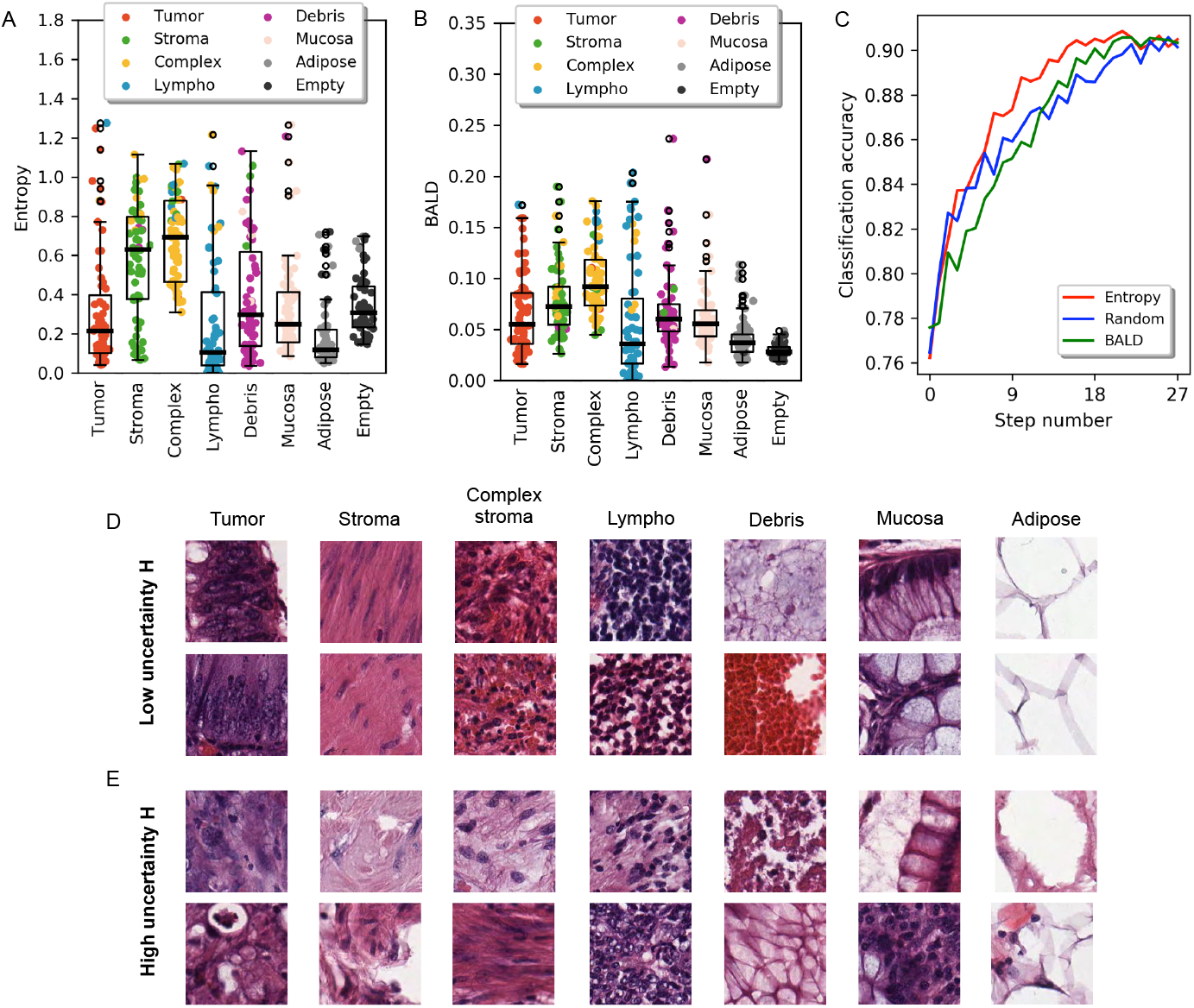
The uncertainty of image classification. **A, B** Distribution of uncertainty for the colorectal cancer images used to train our model. The horizontal axis shows the actual class of these images, whereas the classification of each image is represented with coloured jitter. The y-axis value represents the amount of uncertainty. Our model is on average most uncertain when it comes to the Stroma and Complex classes. It also makes mistakes in classification mostly when it is uncertain. **A** Distribution of uncertainty for the Entropy *H* measure. **B** Distribution of uncertainty for the BALD measure. **C** Results of active learning experiments. Starting from a small training dataset with 320 images in total (step number 0, 40 images per class), the model was re-trained on the dataset increased in every iteration by 160 additional images. Three distinct acquisition functions were tested: Random, Entropy *H* and BALD. At each step, the average classification accuracy was measured (y-axis). **D, E** Microscopic images of tissues composing colorectal cancer. Samples were categorized by uncertainty measured with Entropy *H*. Columns correspond to different tissue classes. **D** Images with low uncertainty *H*. **E** Images with high uncertainty *H*.

### Active learning

Active learning is a set of methods that try to minimise the amount of labelled data needed to fully train a classifier. They start from a small dataset and, as the training goes on, add new training samples according to some kind of acquisition function. We tested the effectiveness of using uncertainty measures as this function, effectively choosing most uncertain images as the ones the model should learn first. The idea here is that if the model learns first what it has the most trouble with, then it should achieve high accuracy at an earlier stage in the active learning process.

Here, we designed an active learning process with either the Entropy *H* or BALD measures acting as acquisition functions (see *Active learning*). We evaluated its efficiency on the *Kather et al.*^29^ dataset by analysing the resulting model accuracy as a function of the number of training samples (Fig. 5C). The Random acquisition function serves as a baseline. In initial active learning iterations the Entropy *H* measure performs very similarly to random selection, but from step 7 (which contained 1440 images) Entropy *H* achieves consistently higher accuracy (with on average 2% improvement in classification accuracy) until the very end of the process. The accuracy of the model trained on samples selected using the BALD measure is worse than the random one from the start of active learning until step 12. From step 13 (which contained 2400 images) it gets slightly better, but never eclipses the accuracy received using the Entropy *H* measure. This proves that the Entropy *H* uncertainty measure can be successfully used as an acquisition function in active learning scenarios utilising our ARA-CNN model. It can speed up the learning process by roughly 45% - the model reaches the classification accuracy equivalent to the full dataset already at step 15, in which the training set contained 2720 images. It means that this subset of images, chosen based on the Entropy *H* uncertainty measure, is large enough to accurately train the model.

### Understanding the uncertainty of image classification

To investigate what pathological features of images are determinant for assigning specific uncertainty values measured by Entropy *H*, we selected test images with very low (*H* ≤ 0.2) and very high (*H* ≥ 0.8) uncertainty and inspected them by eye. We focused on Entopy *H* due to its superior performance in active learning. There were no examples of the Empty class with high uncertainty, indicating this class is easy for the algorithm to recognise and classify properly. For each of the remaining seven tissue classes, images of lowest uncertainty display characteristic pathological features (Fig. 5D). Images of the Tumour class with low *H* display cells that have distinct changes in their nuclei: enlargement, hiperchromasia (dark violet colour), improper chromatin distribution (i.e. spots with higher and lower density) accompanied by multiplication of nucleoli, increased nuclear to cytoplasmic ratio, nuclear fusions and cell overlapping. The images of the Stroma class with lowest uncertainty display typical uniformly stained pink, eosinophilic fibres with elongated nuclei, and low nuclear to cytoplasmic ratio. For the images of the Complex class with low assigned *H*, the stroma is infiltrated by lymphatic or neoplastic cells with addition of erythrocytes. The highly certain images of the Lympho class show features typical for areas of lymphocytic dense infiltration - lymphocytes are intensively stained, monomorphic cells with round nucleus and very scarce thin, basophilic cytoplasm rim. Nucleoli are not visible. Images of the Debris class with low uncertainty *H* values are composed of various tissue samples. First, they contain a mucous, amorphic substance creating multiple, fine vesicles, white in the centre with violet contours. On top of that, features characteristic of the Debris class are mostly extravasated erythrocytes – red, round cell conglomerates presenting very dense collocation with blurred cell contours. Images of the Mucosa class with very low assigned uncertainty show typical features of mucosal glands in large intestine. They are composed of visible characteristic goblet cells that are cylindrical in shape and contain big, round areas filled with mucous - white with violet margin. Small, regular, dark nuclei are visible at the cell periphery. Goblet cells lay in linear or rosette–like formations. Finally, images of the Adipose class with low uncertainty show pathological features typical of the adipose tissue. They are composed of big, white polygonal areas with violet, wide contour, adhering to each other tightly. No nuclei are visible.

In contrast to low uncertainty images, the images with the highest uncertainty show features that are pathologically difficult to categorise (Fig. 5E). For very uncertain images of the Tumour class, the sparse cells visible within the stroma show fewer features of malignancy – most of them are small, regular in shape, with no visible nucleoli. No nuclear fusions or cell overlapping are observed. The pictures could be mistaken with complex stroma. For the images of the Stroma class that were assigned very high uncertainty *H*, the tissue has irregular structure without typical linear fibres and elongated nuclei. Empty spaces in both example images and very low colour intensity in the top one may be artefacts, although whole samples could be categorised as complex stroma or perchance debris because of listed alternations. Out of the two complex stroma example images with very high uncertainty, in the top image (Fig. 5E third column) there are no visible fibres. At the same time, the image contains many pale vesicular areas slightly similar to mucous. The bottom image could be interpreted as a normal stroma sample, because of its colour, fibrotic structure and shape of the nuclei. In the top image representative of very high uncertainty images of the Lympho class, cell arrangement is not very dense and there is a lot of stroma visible between nuclei – this could be categorised as complex stroma instead. The bottom picture shows many features of malignancy that should suggest diagnosis of tumour cells. From the two uncertain example images from the Debris class, the top consists of tissue residues with no particular structures visible. The bottom image shows structures very similar to mucosal glands – areas of mucous are bigger and well margined in comparison to amorphic mucous specific for this category. From the two high uncertainty images of the Mucosa class, the top image has heterogeneous composition. In the right part of the image, goblet cells with their nuclei can be seen. The left part is full of amorphic substance and could be categorised as debris. In the bottom example, only the lower left corner looks like mucosal glands forming rosette. The rest of the image contains stroma with lymphatic infiltration, thus pathologically could be categorised as complex stroma. In the top uncertain example of the Adipose class, although white, empty spaces are clearly visible and cell walls have more irregular margin than normally. In the bottom example, the characteristic polygonal shapes are not visible. The images do not suit any other category more than adipose tissue, however they do not share its typical features.

## Discussion

In this article, we stipulated the necessity of an accurate, reliable and active (ARA) machine learning framework for histopatho-logical image classifiction. We implemented this framework with a new Bayesian deep learning CNN model, called ARA-CNN. ARA-CNN was applied to the task of colorectal tissue classification and incorporated it into an uncertainty-based active pathology workflow. The classification accuracy achieved by our model exceeds the results reported by authors of the training dataset *Kather et al.*^29^ used in this work. The proposed CNN architecture shows outstanding performance in both binary and multiclass classification scenarios, reaching almost perfect accuracy (error rate of 0.89%) in the former case and best in class (error rate of 7.56%) in the latter. It also surpasses the classification performance of other methods that were trained with the same dataset by up to 18.78%.

To achieve reliability, the model measures the uncertainty of each prediction. The benefit of uncertainty assessment is two-fold. First, such a measure can be used to filter out samples that may need a second opinion of a human expert, while the rest (preferably vast majority) can be analysed in a fully automated fashion. Second, as demonstrated by our active learning results (Fig. 5, it can be used to largely reduce the labour that trained pathologists need to put into image labelling and increase the efficiency of model training. In an active learning workflow involving interaction with a pathologist responsible for annotating whole slide images, the pathologist should be informed which classes are the most uncertain and prioritise them in subsequent annotation iterations (Fig. 1A). Our analysis involved a comparison of two different uncertainty measures, Entropy *H*, and BALD. The two measures agreed on which classes are most uncertain on average, pointing to classes which were most often mis-classified by the model. The Entropy *H*, however, outperformed BALD as an acquisition function in the active learning workflow. Compared to random selection, *H* was able to speed-up the training process by a significant margin, while BALD performed only slightly better. Using *H*, the classification accuracy equal to that of the model trained with the full dataset was reached 45% faster. To investigate how the pathological characteristics of images relate to their uncertainty measure *H*, we analysed pathological features of examples of highly certain and highly uncertain images. We observed that highly certain images are very good representatives of their class, while the highly uncertain ones are inconclusive and could have been annotated incorrectly when the dataset was constructed. This shows that measuring uncertainty is a good indicator of how well the model is trained and whether its predictions should be trusted without verification.

The excellent performance of ARA-CNN indicates that it is a step forward in establishing accurate and reliable machine learning models for histopathology. Based on such a model, further exciting avenues of research can be followed. As future work, we plan to apply our model to other histopathological tissue datasets. Due to its deep learning nature, our architecture should easily handle tissue types other than colorectal (potentially with the help of transfer learning). Furthermore, we plan more involved spatial analysis of segmented whole-slide images, especially in conjunction with clinical data. Our segmentation could facilitate application of methods that quantify spatial heterogeneity^56^ in histological samples of colorectal cancer, and improve our understanding of how tumour microenvironment influences the development of this disease. To this end, we plan to work on more precise segmentation algorithms, which will allow better understanding of spatial relations in analysed tissues.

## Data Availability

The model definition is available as open-source Python code on GitHub: https://github.com/animgoeth/pathology-bayesian-cnn

## Acknowledgements

We thank Łukasz Koperski for guidelines in interpreting the histopatological images.

## Author contributions statement

Ł.R. performed all experiments and data analysis and contributed to the model. M.M. developed the model architecture. M.M. and Ł.R. developed the implementation of the approach and prepared the visualisations. J.Z. performed the inspection of images with low and high uncertainty. E.S. supervised the research. M.M. and E.S. conceptualised the project. Ł.R. and E.S. wrote the manuscript. All authors reviewed the manuscript.

